# A photopolymerizable hydrogel enhances intramyocardial vascular cell delivery and promotes post-myocardial infarction healing by polarizing pro-regenerative neutrophils

**DOI:** 10.1101/2022.06.30.497378

**Authors:** Xuechong Hong, Allen Chilun Luo, Ilias Doulamis, Nicholas Oh, Gwang-Bum Im, Pedro J. del Nido, Juan M. Melero-Martin, Ruei-Zeng Lin

## Abstract

The success of vascular progenitor cell transplantation to treat myocardial infarction (MI) is primarily limited by the low engraftment of delivered cells due to a washout effect during myocardium contraction. A clinically applicable biomaterial to improve cell retention is arguably needed to enable optimization of intramyocardial cell delivery. Here, we developed a novel therapeutic cell delivery method for MI treatment based on a photocrosslinkable gelatin methacryloyl (GelMA) hydrogel. A combination of human vascular progenitor cells (endothelial progenitors and mesenchymal stem cells) with the capacity to form functional vasculatures after transplantation, were injected with a rapid in-situ photopolymerization approach into the infarcted zone of mouse hearts. Our approach significantly improved acute cell retention and achieved a long-term beneficial post-MI cardiac healing, including stabilizing cardiac functions, preserving viable myocardium, and preventing cardiac fibrosis. Furthermore, the engrafted vascular cells polarized recruited bone marrow-derived neutrophils toward a non-inflammatory phenotype via TGFβ signaling, establishing a pro-regenerative microenvironment. Depletion of neutrophils canceled the therapeutic benefits produced by cell delivery in the ischemic hearts, indicating that the non-inflammatory, pro-regenerative neutrophils were indispensable mediators of cardiac remodeling. In summary, our novel GelMA hydrogel-based intramyocardial vascular cell delivery approach has the potential to improve the treatment of acute MI.

## 1. Introduction

Myocardial infarction (MI) remains the leading cause of morbidity and mortality worldwide ^1,2^. MI produces irreversible damage to the heart muscle due to a blockage of coronary arteries. Myocardial necrosis starts rapidly after the onset of ischemia, leaving an infarct zone that contains nonfunctional myocytes and excessive inflammatory cells ^3^. MI patients suffer from decreased cardiac outputs, resulting in a downward spiral leading to congestive heart failure ^1^. In the clinics, the acute mortality rate of MI has fallen dramatically through current surgical interventions aided by pharmacotherapy and mechanical devices ^4^. Notwithstanding the short-term benefit of these interventions, only modest improvements in global cardiac functions were observed in patients six months after an acute MI ^5,6^. A significant number of patients go on to develop late sequelae, including heart failure and arrhythmia ^2^. For years, considerable effort has been focused on delivering therapeutic genes, growth factors, and cells to promote post-myocardial infarction healing ^7–11^. However, these strategies are insufficient partly because of the complexity of the hyper-inflammatory microenvironment in the infarcted myocardium ^12–16^.

While the human heart is considered a terminally differentiated organ, adult cardiac regeneration by endogenous repair mechanisms is unfeasible with current medical options ^17^. Over the last decade, many investigators have tested cell-based therapies to prevent adverse ventricular remodeling and promote recovery of MI in animal models, as well as in clinical trials ^8,10,18^. Delivery of therapeutic cells to the heart is performed in two main categories. First, cardiac patches or cell sheets consisting of pluripotent stem cell-derived cardiomyocytes were transplanted to reinforce or replace the injured myocardium aiming to rebuild the loss of heart walls ^19^. These cardiac patches are mainly manufactured by seeding cells in biomaterial scaffolds and preconditioned in a bioreactor before surgical implantation. The second approach injected therapeutic cells directly into the infarcted areas to mitigate the detrimental impact of MI ^20^. In human patients, delivery of cells could be achieved by intramyocardial or intracoronary injection routes through a catheter-based minimally invasive surgical approach, which allows the early therapeutic intervention in the acute MI phase ^9^. Examples of tested adult stem/progenitor cells include cardiosphere-derived cells ^21^, bone marrow-derived mononuclear cells ^22,23^, endothelial progenitor cells ^24^, and mesenchymal stem cells ^25^. However, except for cardiomyocytes, the long-term engraftment of injected cells was shown to be unnecessary to exhibit therapeutic effects. Accumulating evidence suggests non-myocyte cell types can provide beneficial effects through paracrine factors aiding cardiac remodeling or the modulation of inflammatory response ^7,12,25^. However, the underlying mechanisms are not well-understood.

Delivering vascular progenitor cells is a promising new therapeutic strategy for ischemic cardiomyopathy. Some studies have focused on delivering vascular cells to build new blood vessels and stimulate angiogenesis ^26–28^. The concept was to enable rapid restoration of microvascular networks within the ischemic myocardium to mitigate adverse remodeling. For instance, in a rat myocardial ischemia/reperfusion injury (IRI) model, animals injected with human endothelial colony-forming cells (ECFCs) and mesenchymal stem cells (MSCs) in phosphate-buffered saline (PBS) revealed a modest reduction in adverse ventricular remodeling ^26^. The quantitative analysis showed a rapid loss of human cells shortly after myocardial injection, suggesting the lack of sufficient cellular retention could be the major reason for the weak therapeutic effect ^26^. Therefore, it is still an open question whether human vascular progenitor cells with vasculogenic capability can mitigate adverse remodeling and promote functional recovery.

Low cell retention and engraftment are the major obstacles to achieving a significant functional benefit in MI, irrespective of the cell type used ^29^. Beating hearts actively push therapeutic cells injected in liquid vehicles out of the myocardium, resulting in low acute cell retention between 0.5-10% across several studies with different cell types, animal models, and delivery routes ^29–35^. To address this problem, several biomaterials for enhancing cell engraftment have been tested in the last decade ^36–39^. In this study, our approach is to use well-characterized Gelatin methacryloyl (GelMA) hydrogel to deliver human ECFCs and MSCs to build neo-vascular networks in the ischemic myocardium. GelMA was synthesized in our previous studies to be a photocrosslinkable, bioadhesive hydrogel following its injection in tissues ^40–42^. Cell-laden GelMA formulation can polymerize rapidly (10 s upon exposure to UV light in the presence of a photoinitiator) in myocardial tissues without compromising the therapeutic cell viability. We proposed that this rapid polymerization would be a critical feature to avoid hydrogel dissemination at the implantation site. Recently, we also demonstrated that this material is fully compatible with ECFC-based vascular morphogenesis ^41^, and thus we proposed its use for vascular cell therapy in infarct hearts. Here, we demonstrated that GelMA hydrogel can be transmyocardially photocrosslinked and maintain vascular cell retention in infarcted mouse hearts. GelMA-enabled cell delivery promoted infarct healing and adaptive remodeling to preserve cardiac function. Importantly, enhanced cell retention allows us to study and discover the early beneficial effects of human vascular cells in MI for the first time. We found that the enhanced retention of human ECFCs and MSCs enabled the perivascular-endothelial cell interactions to upregulate transforming growth factor beta (TGFβ) expression. TGFβ signaling promoted the polarization of non-inflammatory, pro-regenerative neutrophils, which were shown to be indispensable mediators of beneficial cardiac remodeling.

## 2. Results

### 2.1 In situ polymerization of GelMA hydrogel for cell delivery in infarcted myocardium

Transmyocardial polymerization of GelMA for cell delivery was evaluated in vivo using a mouse MI model (**Figure 1**). MI was induced by permanent ligation of the left anterior descending (LAD) arteries (**Figure S1A** and **Video S1**, Supporting Information) in 10-12-week-old SCID mice. A mixture of human vascular progenitor cells (ECFCs and MSCs; 5×10^5^ for each cell type) suspended in 5% (w/v in PBS) GelMA precursor solution was injected into the infarcted myocardium within 10 min of LAD ligation. The total injection volume per heart was 100 μL. We first demonstrated that this injection volume was sufficient to cover the infarct zone (**Figure S1B**). We then determined that UV light could transmit through the myocardium of mice using our OmniCure S2000 UV lamp (wavelength 320-500 nm; the intensity of 40 mW/cm^2^). GelMA polymerization was achieved after just 10 seconds of transmyocardial exposure to UV light (**Video S2**). The general safety of short UV illumination by our UV lamp had been evaluated in a previous study ^42^. Our working range of UV light exposure (10 - 45 seconds) was deemed safe for both injected cells and surrounding tissues ^42^. Moreover, we did not observe leakage of GelMA solution or swelling of local tissues with the injection volume and crosslinking conditions used.

**Figure 1.**
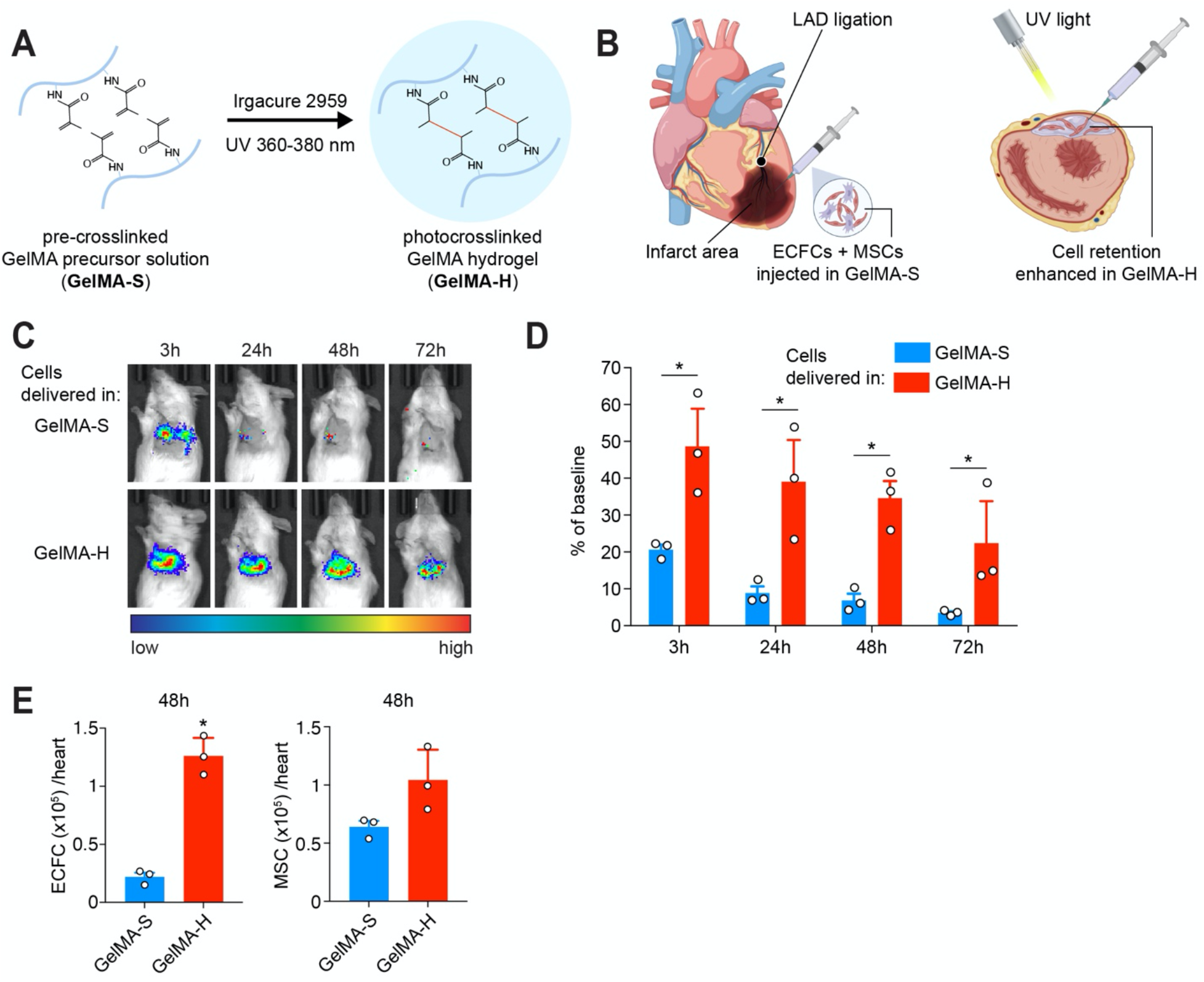
Enhanced intramyocardial retention of human vascular progenitor cells delivered in a photocrosslinked GelMA hydrogel. (A) Schematic depicting the photocrosslinking of GelMA from a precursor solution (GelMA-S) to a hydrogel matrix (GelMA-H) under UV irradiation. (B) Schematic depicting the intramyocardial cell injection. MI was surgically induced by LAD ligation. Human vascular progenitor cells (ECFCs + MSCs) resuspended in GelMA precursor solution were injected into three locations in the infarcted area. UV light was immediately applied on top of the infarcted area to polymerize the cell-laden hydrogel (referred to as GelMA-H group). UV irradiation was omitted for the control group to maintain cells injected in GelMA precursor solution (GelMA-S group). (C) Viable cell retention was measured by bioluminescence imaging of human ECFCs expressing a luciferase reporter. (D) Total bioluminescent signals in each animal’s chest region were measured on 3, 24, 48, and 72 h post-MI and compared to the baseline (i.e., cells delivered to arrested mouse hearts). (E) Retention of human ECFCs or MSCs was quantified by flow cytometric analysis on 48 h post-MI. Data correspond to cell numbers from each harvested MI heart. Data are mean ± s.d.; n = 3 mice per group (indicated by individual dots). \**P* < 0.05 between GelMA-H and GelMA-S. Statistical methods: two-way ANOVA followed by Bonferroni’s post-test analysis (D) and unpaired two-tailed Student’s t-tests (E).

### 2.2 GelMA hydrogel enhanced cell retention in ischemic hearts

To evaluate the acute myocardial retention of human vascular progenitor cells, we injected 5×10^5^ luciferase-labeled ECFCs (luc-ECFCs) with 5×10^5^ non-labeled MSCs per heart and quantified the bioluminescence intensity in the infarcted myocardium (**Figure 1C**). We compared transmyocardial photocrosslinking of cell-laden GelMA hydrogel following injection into the infarct zone (referred to as GelMA-Hydrogel or **GelMA-H** group; **Figure 1B**) with the results of cell injection in the same GelMA precursor solution but without UV illumination (GelMA-Solution or **GelMA-S** group). Notably, the GelMA precursor solution remained liquid in physiological conditions (i.e., at 37°C and in contact with blood or tissues) if not crosslinked by UV light ^43^. Since cardiac contraction is primarily responsible for the early washout of cells from the injection site ^31^, we injected the same number of human cells into arrested mouse hearts and performed a bioluminescence measurement after 15 min to obtain a baseline. Thus, bioluminescence signals compared to this baseline were used to estimate the percentage of ECFC retention (**Figure 1D**).

As early as 3 h after MI induction and cell delivery, the presence of ECFCs in the GelMA-S group was rapidly reduced to 20.6 + 2.3% (**Figure 1C-D**), a low retention rate similar to previous studies describing intramyocardial cell delivery in PBS ^22,31,44^. Cardiac contraction potentially reduces cell retention by extrusion of the liquid injectate during each heartbeat. Therefore, photopolymerization of GelMA hydrogel should, in principle, minimize this cell backwash. Indeed, in the GelMA-H group, ECFC retention was significantly improved to 48.6 + 13.6% (3 h, \**P* < 0.05 vs. cells in GelMA-S; **Figure 1D**), revealing a dramatic effect of sealing the injection site on cell retention. The higher ECFC retention in the GelMA-H group was maintained throughout the 3-day monitoring post-MI (**Figure 1D)**.

To confirm that the GelMA-H enhanced the retention of both endothelial and mesenchymal cells, we quantified non-labeled ECFCs and MSCs in infarcted hearts by flow cytometry (**Figure S2**, Supporting Information). Cell delivery in GelMA-H significantly enhanced both ECFC and MSC retentions at 48 h compared to the GelMA-S group (**Figure 1E**; *P* value = 0.0005 for ECFCs and 0.0728 for MSCs). Compared to the number of injected cells, the GelMA-H group preserved 25.2% of viable ECFCs and 20.8% of MSCs, while the GelMA-S group only retained 4.4% of ECFCs and 12.8% of MSCs. Notably, GelMA hydrogel maintained the engraftment of ECFCs and MSCs close to the original 1:1 ratio, which was optimized to achieve long-term vascularization ^45^. Collectively, our data suggest that the rapid in situ photopolymerization of GelMA hydrogel adequately protected the human vascular progenitor cells and reduced the “washout” effect during the acute phase (72 h) following MI.

### 2.3 Human vascular progenitor cells delivered in GelMA hydrogel improve cardiac remodeling

To determine whether the improvement in acute cell retention would impact post-MI cardiac healing, we used echocardiography to evaluate the cardiac functions: left ventricular fractional shortening (LVFS) and left ventricular ejection fraction (LVEF) (**Figure 2 and S3**, Supporting Information). Echocardiography revealed no significant difference in baseline cardiac function between groups measured 1 h before LAD ligation. On day 7 following the induction of MI, LVFS and LVEF were higher in the mice treated with human cells delivered in GelMA-H compared to the untreated or GelMA-S-treated animals (**Figure 2B-C;** 30.3 + 2.9%, 24.6 + 3.2%, 25.2 + 2.8% for LVFS; 66.4 + 4.5, 56.9 + 5.6, 56.8 + 4.6 for LVEF, respectively). After treatment, both cardiac functions were significantly higher in the GelMA-H group. Compared with the GelMA-S group, LVEF and LVFS in the GelMA-H group were improved considerably on days 7, 14, and 21 (increased 20.2%, 18.7%, and 19.7% for LVFS; 16.9%, 17.4%, and 17.1% for LVEF). Both untreated and GelMA-S-treated groups showed a continuous decline in LVEF and LVFS from day 3 to 28 post-MI (MI-untreated: reduced 17.6% for LVFS and 13.4% for LVEF; GelMA-S: reduced 19.6% for LVFS and 14.3% for LVEF), indicating a deterioration of cardiac function due to adverse infarct remodeling. There was no significant difference in LVEF and LVFS between untreated or GelMA-S-treated groups, suggesting a lack of therapeutic benefit in groups with low cell retention. In contrast, in the GelMA-H group, cardiac function was generally preserved from day 3 to day 28 (GelMA-H: reduced only 9.5% for LVFS and 6.7% for LVEF).

**Figure 2.**
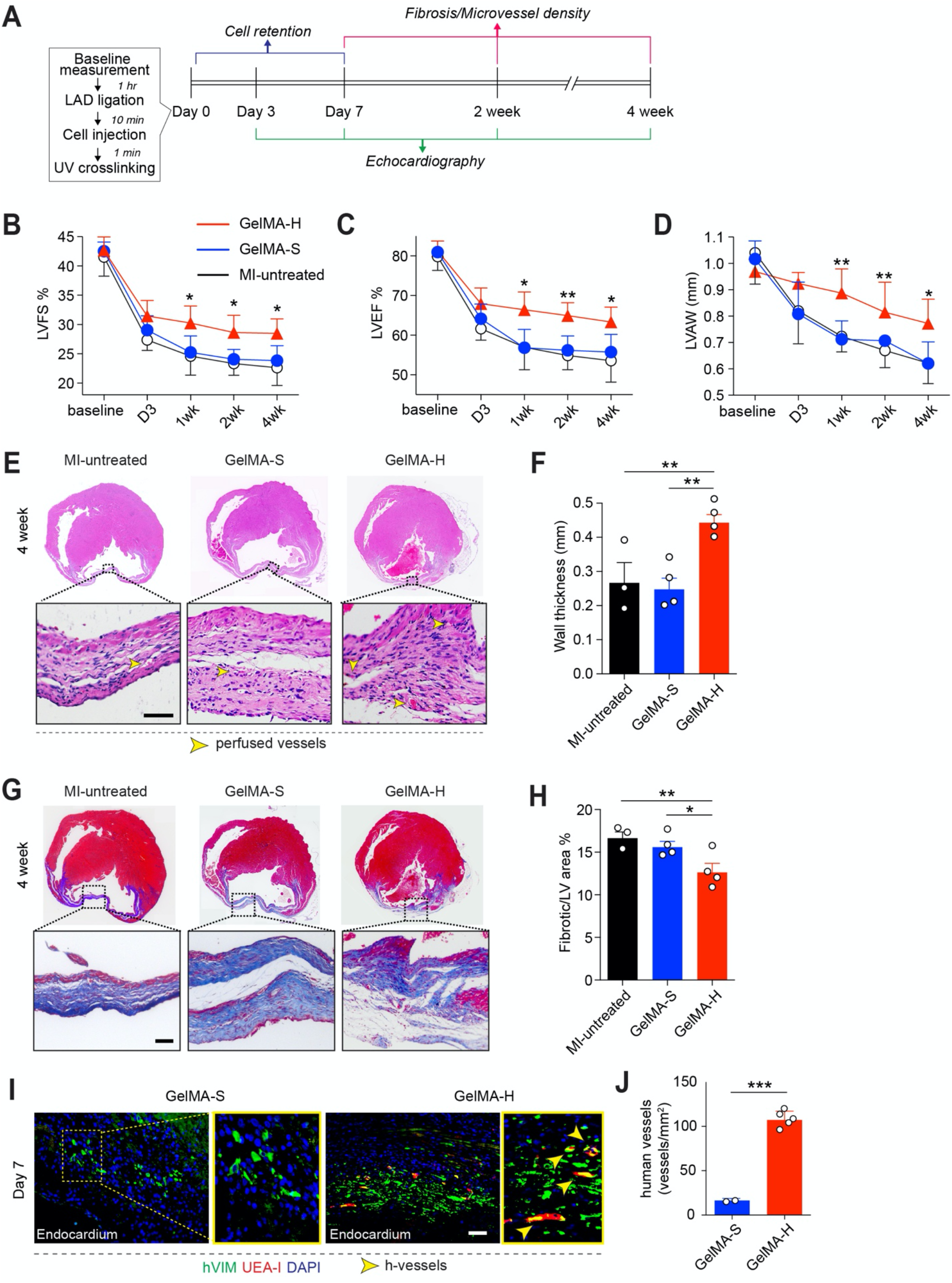
Human vascular progenitor cells delivered in GelMA hydrogel improved cardiac remodeling. (A) Diagram of experimental schedule. (B-D) Echocardiography analysis for cardiac function. Echocardiography was performed on anesthetized mice at 3 days and 1, 2, and 4 weeks post-MI following cell delivery in GelMA-H or GelMA-S (n=6 for each group). Mice received LAD ligation, but no cell injection (MI-untreated) served as control (n=6). Baseline data were obtained from mice within 1h before performing LAD ligation. (B) Left ventricular fractional shortening (LVFS). (C) Left ventricular ejection fraction (LVEF). (D) Left ventricular anterior wall thickness (LVAW). (E) Representative heart horizontal panoramic views (upper) and microscopic images (lower) of myocardial sections stained with H&E at 4 weeks post-MI. Scale bars, 100 μm. (F) The LV wall thickness in the infarcted areas quantified from H&E sections. (G) Representative heart sections stained with Masson’s trichrome at 4 weeks post-MI. Scale bars, 100 μm. (H) The percentage of fibrotic areas in the total LV cross-sections quantified by Masson’s trichrome staining. (I) Histological analysis of the infarcted myocardium on day 7 revealed the presence of perfused human blood vessels only when cells were delivered in GelMA-H group, but not in GelMA-S. Human microvessels were identified by immunostaining for human-specific vimentin (hVIM) and UEA-I lectin binding. Scale bars, 100 μm. (J) Total perfused human microvessel density quantified in the infarcted myocardium. Statistical methods: ANOVA followed by Bonferroni’s post-test analysis (B-D) and unpaired two-tailed Student’s t-tests (F, H, J). Data present mean + s.d. *\*P* < 0.1, *\*\*P* < 0.05, \*\*\**P* < 0.001.

Myocardium salvage after MI is the single most important determinant factor for the long-term preservation of cardiac function and survival ^2^. We measured the myocardial wall thickness by echocardiography. The GelMA-H group exhibited a significantly higher left ventricular anterior wall thickness (LVAW) compared to the untreated or GelMA-S groups (**Figure 2D)**. Again, no improvement was found by comparing GelMA-S and untreated groups, suggesting that high cell retention was required for beneficial cardiac remodeling. The preservation of the myocardium was confirmed by histological analysis of cross-sections in the infarct zone after 4 weeks (**Figure 2E**). In mice that received cells in GelMA-H, myocardial thickness significantly improved compared to the untreated or GelMA-S groups (**Figure 2F;** 0.44 + 0.05 mm vs. 0.25 + 0.06 mm, \*\**p* < 0.05). Masson’s trichrome staining of cardiac tissues harvested at 4 weeks also confirmed that the GelMA-H group showed significantly reduced myocardial fibrosis compared to the untreated or GelMA-S-treated groups (**Figure 2G-H**). Taken together, our data indicate that improving acute cell retention translates to beneficial cardiac remodeling, including enhanced revascularization, less scar tissue formation, and stabilization of overall cardiac functions after MI.

### 2.4 Functional human vasculature in infarct hearts

We sought to investigate the therapeutic mechanism resulting from the delivery of human vascular progenitor cells in GelMA-H. First, we examined the infarcted hearts on day 7 to assess the formation of human blood vessels ^26^. The histological examination carried out by immunofluorescence staining of human-specific vimentin antibodies (stain both human ECFCs and MSCs) and UEA-I lectins (bind specifically to human ECFCs, but not mouse endothelial cells) revealed that the extent of vascular network formation differed in the GelMA-H and GelMA-S groups (**Figure 2I**). Significant engraftment of human ECFCs and MSCs in the GelMA-H groups was evidenced by the formation of human-specific vascular networks inside the infarcted myocardium (**Figure 2J**). In the GelMA-S group, only a few human MSCs, but not ECFCs, were stained in the infarcted myocardium. These data suggested that the in situ photocrosslinking of the GelMA hydrogel significantly improved the retention of co-delivered human ECFCs and MSCs. Without the timely gelation of GelMA hydrogel, although some MSCs could engraft, ECFCs failed to participate in the revascularization of the infarcted myocardium.

The formation of functional human vasculature in ischemic tissues might explain parts of the therapeutic benefit observed in the GelMA-H group. However, we noticed significant improvement of LVAW in the GelMA-H group (**Figure 2D**) as early as day 3, and improved cardiac functions by day 7 (LVFS and LVEF; **Figure 2B-C**). Considering that the onset of human blood vessel perfusion takes approximately 7 days in this model of vascular progenitor cell implantation ^41,42,46– 48^, this timing inconsistency between human cell-mediated revascularization and beneficial cardiac remodeling suggested that other therapeutic pathways might be involved in preserving cardiac functions during the early stage of MI.

### 2.5 Gene expression analysis upon human vascular progenitor cell delivery in infarcted hearts

To further elucidate the molecular aspects of the therapeutic benefits produced by ECFC+MSC delivery in GelMA-H, we conducted an exploratory transcriptional analysis using RNA sequencing (RNAseq). The infarcted left ventricle tissues were harvested from GelMA-H and GelMA-S groups on day 2 post-MI to study the change of gene expression profiles in the early stage of treatments. Tissues from uninjured hearts served as controls.

Globally, we detected differentially expressed genes (DEGs) among the three analyzed groups (**Figure S4A**). Hierarchical clustering analysis (**Figure S4B**) revealed the proximity between two post-MI groups (GelMA-H and GelMA-S) in comparison to uninjured controls. This pattern was confirmed by the principal component analysis (PCA; **Figure S4C)**. We performed a Gene Ontology (GO) analysis of the DEGs between post-MI GelMA-H and uninjured controls (**Figure S4D**). The results showed that the top enriched GO categories due to infarct injury and therapeutic cell delivery were associated with positive regulations of immunity, including inflammatory response, immune response, and leukocyte chemotaxis. These findings suggest that the immune system was deeply involved in the early stage of the MI process.

To gain more insight into the transcriptional difference between GelMA-H and GelMA-S groups, we carried out a pairwise analysis of DEGs. There were 137 genes significantly upregulated in the GelMA-H group whereas 617 genes were downregulated (**Figure S4A**, GelMA-H vs. GelMA-S). The volcano plot (**Figure S4E**) shows the up- and down-regulated genes comparing the GelMA-H group to the GelMA-S group. Notably, we observed upregulation of several genes related to neutrophil development and maturation, including expression of *s100a8, s100a9, elane, ngp, mpo, camp, lef1, and ms4a2* in the GelMA-H group. This was associated with upregulation of genes implicated in immature neutrophil trafficking (*sdf, cxcr4*) and hematopoiesis (*gata1, hemgn*) in GelMA-H vs. GelMA-S (**Figure S4F-G**). Altogether, the data suggested a potential effect of human ECFCs+MSCs in GelMA-H on the recruitment and polarization of neutrophil subpopulations in the early stage of MI, although it is important to note that the data variation between samples within the GelMA-H group was high.

### 2.6 Recruitment of neutrophils with reparative phenotype in GelMA-H group

RNAseq data analysis suggested the difference in neutrophil-related genes between GelMA-H and GelMA-S groups. Previously, we demonstrated that in vivo implanted human vascular progenitor cells could alternatively polarize neutrophils into a non-inflammatory phenotype, mediating vascular assembly and anastomosis ^47^. Moreover, recent studies have shown that pro-regenerative neutrophils (herein termed NR; “R” for regeneration) were implicated in orchestrating tissue repair in various organs ^49–59^. However, the role of NR in post-MI healing remains unclear. Therefore, we focused our studies on the polarization of the neutrophils following MI.

We first examined the early-stage host cell infiltration post-MI (**Figure 3**). We analyzed by flow cytometry the recruited mouse CD45^+^ leucocytes on day 2 in infarcted myocardium and classified them into Ly6G^+^F4/80^-^ neutrophils or Ly6G^-^F4/80^+^ macrophages (**Figure 3A**). Infiltration of Ly6G^-^F4/80^-^ lymphocytes was negligible, as expected in immunodeficient mice ^47^. Also, the presence of murine resident myeloid cells in the uninjured myocardium was low (**Figure 3B)**. LAD ligation followed by either GelMA-H or GelMA-S treatment significantly recruited host myeloid cells (**Figure 3B)**, which is consistent with the induced innate immune response observed by RNAseq analysis. Moreover, the GelMA-H group recruited more neutrophils and macrophages than the GelMA-S group (**Figure 3B)**, which is consistent with our previous observation on the engagement of host myeloid cells by ECFCs and MSCs ^47^. We further evaluated neutrophil subpopulations using flow cytometry and classified Ly6G^+^F4/80^-^ cells into naïve neutrophils (N0; CXCR2-), pro-inflammatory neutrophils (NI; CXCR2^+^CD206^-^), and pro-regenerative neutrophils (NR; CXCR2^+^CD206^+^; **Figure 3A and C**). The NI was the dominant subpopulation (83.27% of total neutrophils; **Figure 3D**) in MI mice treated with GelMA-S, which is consistent with previous report that showed pro-inflammatory neutrophils significantly mediating adverse cardiac remodeling in the first 7 days post-MI ^52^. In contrast, NIs were considerably reduced (18.02%) in the GelMA-H group, where NRs were significantly more abundant (57.66% of total neutrophils), suggesting a shift toward NR polarization. The GelMA-H group also recruited more naïve neutrophils from the bone marrow than that in the GelMA-S group (24.32% vs. 13.31%). Together, these data suggested that improving human vascular progenitor cell retention could translate into a pro-regenerative host neutrophil polarization during the early stage of MI healing. Since NRs have been previously shown to promote tissue healing by facilitating the resolution of inflammation and stimulating angiogenesis, cell proliferation, and ECM remodeling ^47,60–62^, the increased abundance of NR in the GelMA-H group could in part explain the beneficial effects observed in cardiac function.

**Figure 3.**
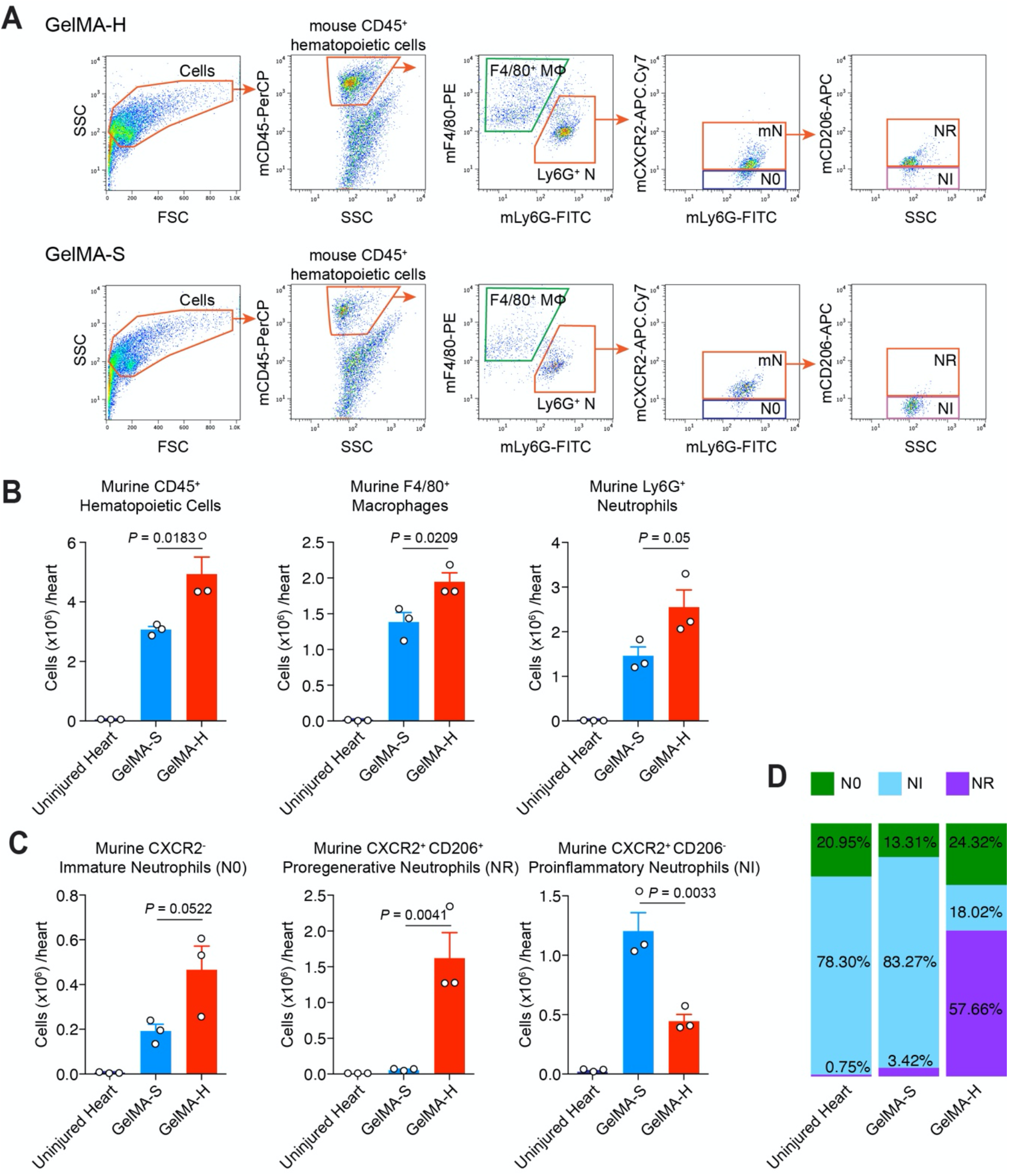
Human vascular progenitor cell retention promotes the recruitment of neutrophils with a reparative phenotype. The infarcted myocardial tissue was harvested on day 2 post-MI. Tissues were digested into single-cell suspensions for flow cytometric analysis. (A) Flow cytometry gating strategy to identify recruited myeloid cells. Representative flow cytometric analyses of ischemic myocardium injected with ECFC+MSC in GelMA-S or GelMA-H. MΦ, macrophages; N, neutrophils; mN, mature neutrophils; N0, naïve neutrophils; NR, pro-regenerative neutrophils; NI, pro-inflammatory neutrophils. (B) Quantitative cytometric analyses of murine CD45+ hematopoietic cells, Ly6G^-^F4/80^+^ macrophages, and total Ly6G^+^F4/80^-^ neutrophils obtained from the ischemic myocardium. The myocardium from uninjured hearts served as controls. (C) Recruited Ly6G^+^F4/80^-^ neutrophils were classified into three subpopulations: N0 (CXCR2^-^), NI (CXCR2^+^CD206^-^), and NR (CXCR2^+^CD206^+^). (B,C) Data are mean ± s.d.; n = 3 per group; *P* values between GelMA-S and GelMA-H are listed. (D) Proportions of the three neutrophil subpopulations in each group.

### 2.7 Human vascular cell interaction upregulated NR polarization factor TGFβ1

We next examined if the prolonged retention and interaction between human ECFCs and MSCs in the GelMA-H group resulted in a release of paracrine factors that could promote NR polarization and whether these factors were absent in the GelMA-S group. To this end, we collected conditioned media from ECFC+MSC coculture in GelMA hydrogel in vitro (CM^CO^; **Figure 4A**). Next, murine naïve neutrophils isolated from bone marrow (BM) were cultured in CM^CO^ for 24 h. NR polarization was evidenced by the upregulation of *Il4, Vegfa* and *Arg1* gene expressions upon exposure to CM^CO^ (**Figure 4B**). However, the evidence for NR polarization was minimal when BM neutrophils were cultured in basal medium or conditioned media obtained from ECFC or MSC monoculture (CM^ECFC^ or CM^MSC^), suggesting an important role for ECFC-MSC crosstalk in the polarization of NR.

**Figure 4.**
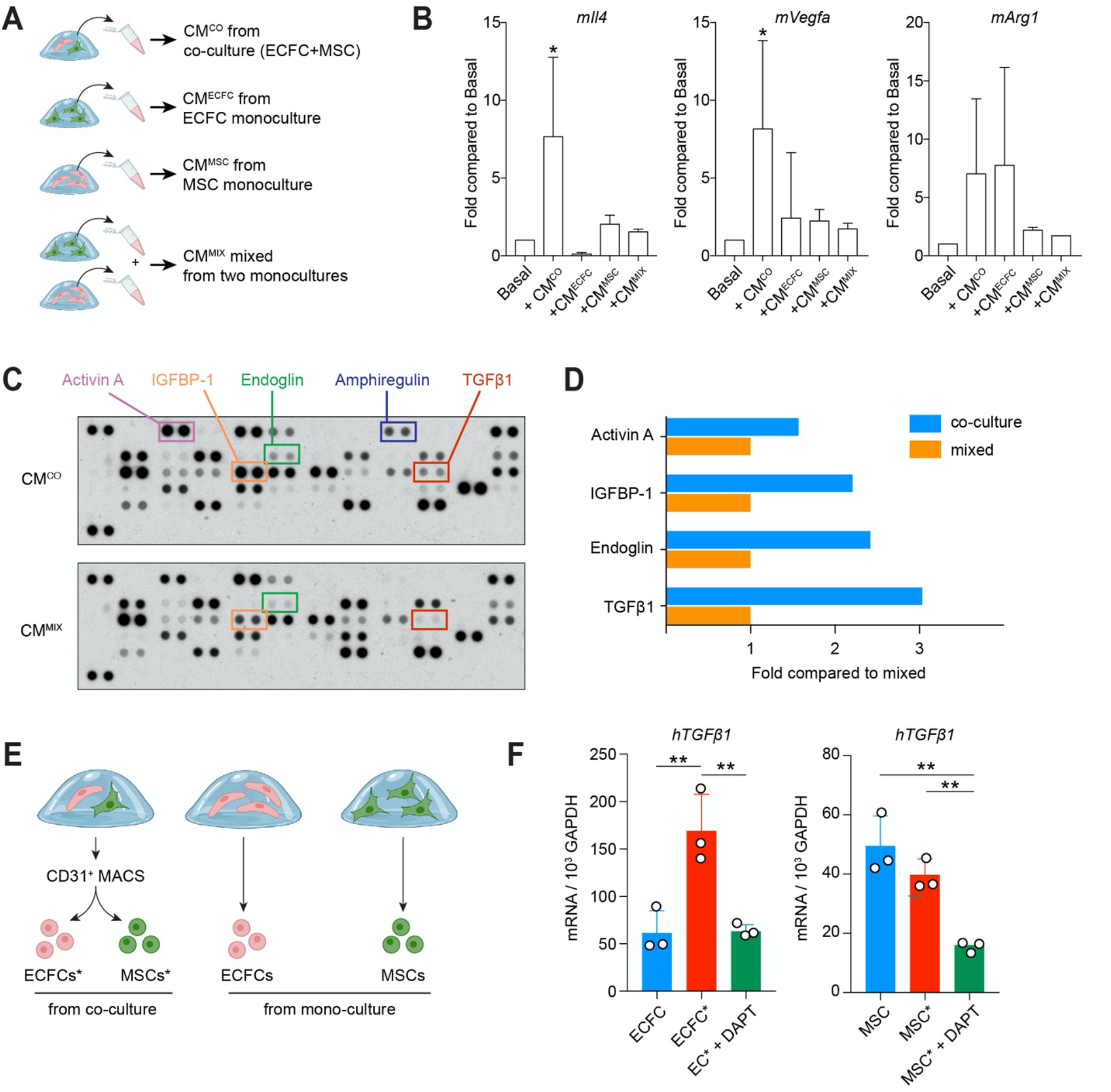
Human vascular progenitor cell crosstalk upregulates NR polarization factor TGFβ1. (A) Conditioned media (CM) were collected from ECFC+MSC coculture in GelMA hydrogels in vitro (CM^CO^). CMs obtained from ECFC or MSC monoculture (CM^ECFC^ or CM^MSC^) and mixed in 1:1 ratio (termed CM^MIX^) served as a control. (B) Mouse naïve neutrophils isolated from bone marrow were cultured in basal medium or indicated conditional media for 24 h. Gene expressions of *Il4, Vegfa*, and *Arg1* expressions in mouse naïve neutrophils were measured by the quantitative PCR analyses and compared to the levels in basal medium. Data are mean ± s.d.; n = 3 per group. \**P* < 0.05 between Basal and CM^co^ groups. (C) The secretions of cytokines in conditioned media were measured by proteomic arrays. Selected cytokines are marked with colored outlines. (D) Quantification of cytokine levels was carried out by densitometry. (E) ECFC+MSC were cocultured in GelMA hydrogel and separated by magnetic-activated cell sorting (MACS) for gene expression analyses. ECFC or MSC monoculture served as controls. (F) Quantitative PCR analyses of TGFβ1 expression in ECFCs sorted from ECFC+MSC coculture (labeled ECFC*) or from ECFC monoculture (ECFC). The same labeling applied to MSC (monoculture) and MSC* (sorted from coculture). DAPT was added to ECFC+MSC coculture to block Notch-mediated perivascular interaction. Data are mean ± s.d.; n = 3 per group; \*\**P* < 0.01 between indicated groups.

To gain further insight into the NR polarization factors, we performed a proteomic analysis of the conditioned media from the ECFC+MSC coculture (**Figure 4C-D**). Mixed conditioned media (1:1 mixture of CM^ECFC^+CM^MSC^; termed CM^MIX^) obtained from separate ECFC and MSC monocultures served as a control. The secretion of several growth factors was induced in the ECFC+MSC coculture, including Activin A, IGFBP-1, Endoglin, Amphiregulin, and TGFβ1. The increase in TGF-β1 was consistent with our previous study that showed TGFβ1 promotes NR polarization ^47^. We confirmed at the mRNA level that the ECFC+MSC coculture upregulated TGFβ1 expression only in ECFCs but not in MSCs (**Figure 4E-F**). Moreover, exposing the ECFC+MSC coculture to the Notch inhibitor DAPT (a γ-secretase inhibitor) blocked the induction of TGFβ1 expression (Notch signaling is known to mediate vascular cell interaction ^47^). Together, these results suggested that NR polarization was directly related to the engrafted human vascular progenitor cells in the infarcted tissues. Notch-mediated interaction of ECFCs and MSCs could modulate the early innate immune landscape by increasing TGFβ1 secretion by the ECFCs.

### 2.8 Neutrophil depletion abrogates the improvement in cardiac remodeling

To confirm the pivotal role of host neutrophils during cardiac remodeling, we depleted circulating neutrophils in MI mice treated with GelMA-H. To this end, we treated mice from two days before LAD ligation to post-operative day 6 with an anti-Gr1 or IgG (200 μg) control antibody given every 2 days via intraperitoneal injection (**Figure 5A**). This treatment with anti-Gr1 antibody was effective at depleting circulating neutrophils without altering blood monocytes and lymphocytes (**Figure 5B-C**). Importantly, neutrophil depletion abolished the improved post-MI cardiac functions observed at days 3 and 7 in the GelMA-H group (LVEF and LVFS; **Figure 5D, 5E**) and significantly reduced myocardium thickness at day 7 (LVAW, **Figure 5F**). All three measurements of cardiac functions (LVEF, LVFS, and LVAW) in neutrophil-depleted mice were comparable to those observed in the untreated or GelMA-S groups (**Figure 2**). Together, these findings suggest that depletion of neutrophils leads to excessive loss of myocardium that might explain the progressive worsening of cardiac function. Indeed, host neutrophils were indispensable for the therapeutic effects exhibited by human vascular cells delivered in GelMA-H.

**Figure 5.**
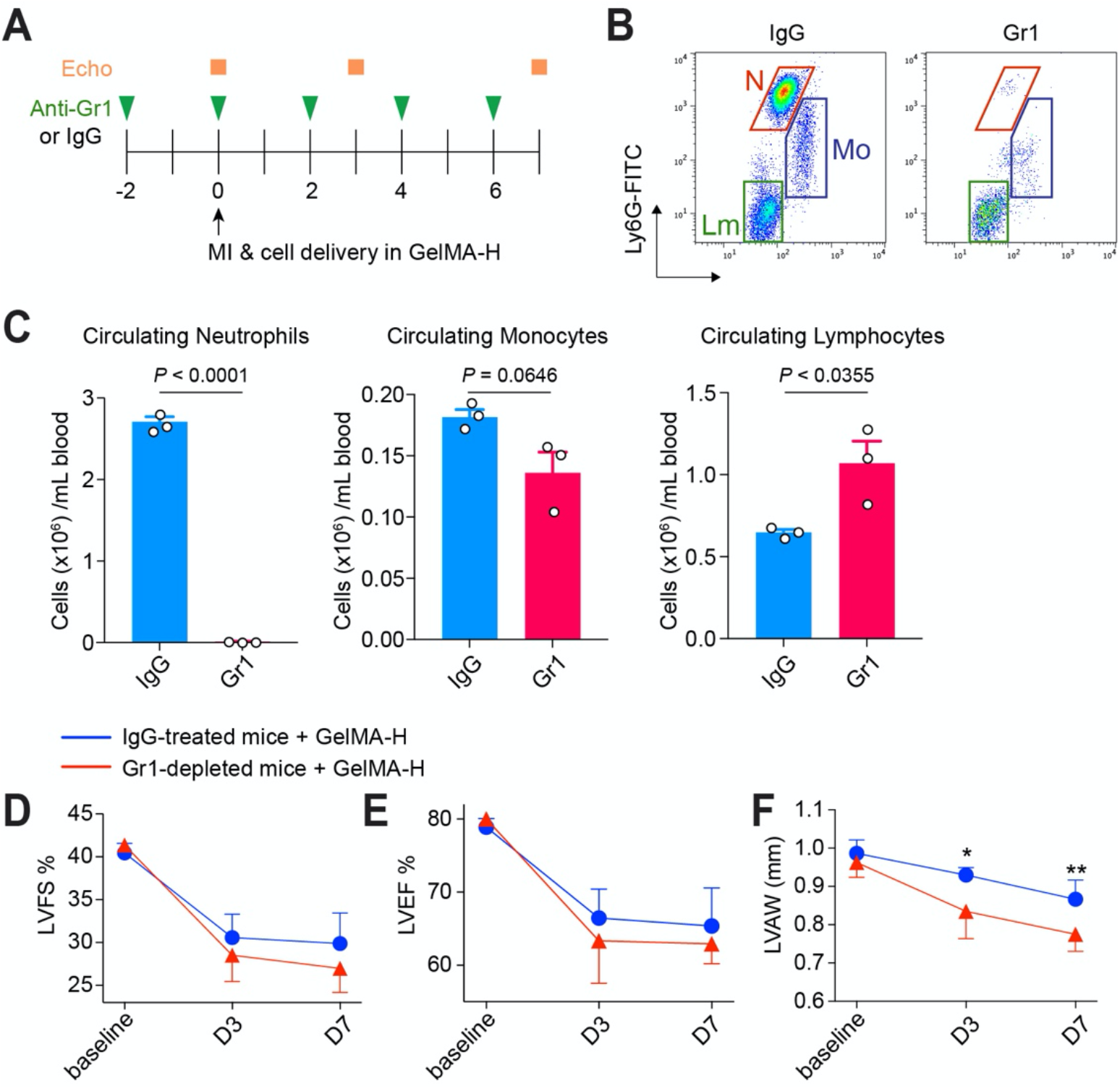
Neutrophil depletion abrogated the improvement in cardiac remodeling of human vascular cell delivery. (A) Experimental schedule: anti-Gr-1 or IgG control antibody (200 μg each) was given via an intraperitoneal injection every 2 days. ECFC+MSC were delivered in GelMA-H into the ischemic myocardium at day 0. The effect of anti-Gr-1 depletion was monitored on days 3 and 7 post-MI by echocardiography. (B) Representative flow cytometry analyses of circulating leukocytes from the blood of anti-Gr-1-treated or IgG-treated mice on day 2. N, neutrophils; Mo, monocytes; Lm, lymphocytes. (C) Quantitative cytometric analyses of circulating leukocytes in the peripheral blood of anti-Gr-1-treated or IgG-treated mice. Data are mean ± s.d.; n = 3 per group; *P* values between IgG and Gr1 treatments are listed. (D-F) Echocardiography analysis for cardiac function. Echocardiography was performed on anesthetized mice at days 3 and 7 post-MI following cell delivery in GelMA-H. Baseline data were obtained from mice within 1h before performing LAD ligation. (D) Left ventricular fractional shortening (LVFS). (E) Left ventricular ejection fraction (LVEF). (F) Left ventricular anterior wall thickness (LVAW). Data are mean ± s.d.; n=3 for IgG-treated and n=4 for Gr1-treated. *\*P* < 0.1, *\*\*P* < 0.05.

## 3. Discussion

An effective early intervention to achieve sufficient cell retention in the infarcted myocardium has important implications for cardiac remodeling. In the present study, we demonstrated that human vascular progenitor cell retention in MI hearts could be significantly improved by delivering the cells in a photocrosslinked GelMA hydrogel. In contrast, cells injected in GelMA precursor solution were rapidly lost from the beating hearts. The improved acute cell retention showed a significant impact on post-MI cardiac remodeling. Echocardiography and histological examination demonstrated that the delivery of ECFC+MSC in the GelMA hydrogel prevented a drastic loss of cardiac functions, preserving viable myocardium, and averting cardiac fibrosis. The delivered human ECFCs and MSCs formed perfused vascular networks in the ischemic microenvironment. However, vascular regeneration was not the sole mechanism leading to cardiac healing. The human vascular cells also mediated the pro-regenerative polarization of recruited neutrophils, which in turn facilitated cardiac remodeling.

Leukocyte infiltration in ischemic areas is a hallmark of MI. The inflammatory response appears a double-edged sword because it is a protective attempt by the organism to initiate the healing process; however, prolonged inflammation can exacerbate scarring and loss of organ function ^15,63^. Overwhelming infiltration of innate immune cells has been shown to promote adverse remodeling and cardiac rupture ^64^. The successful resolution of inflammation appears as a critical event in the repair of tissue damage, but little is known about the primary mediators of resolution in the infarcted cardiac tissue ^65–67^. Neutrophils massively infiltrate the heart soon after MI, where they can promote both tissue repair and damage ^14,49^. Emerging evidence indicates that neutrophils comprise two distinct subpopulations with different polarized phenotypes: a canonical pro-inflammatory phenotype (referred to as NI) and an alternative anti-inflammatory, pro-regenerative, phenotype (NR), resembling the well-established ‘M1–M2’ macrophage paradigm ^58,68,69^. Previous reports have suggested that cardiac neutrophils follow a temporal polarization pattern after MI, switching from a pro-inflammatory NI to an anti-inflammatory NR profile ^13,50,52,70,71^. However, this switch could take 5-7 days post-MI in rodent models, while excessive inflammatory cells had produced irreversible damage to the surrounding myocardium ^52^. Here we showed that an optimized delivery of human vascular progenitor cells could promote the alternative polarization of recruited host neutrophils in the ischemic tissues as early as 2 days post-MI. NRs have been shown to actively contribute to the repair of various tissues, mainly through suppressing pro-inflammatory reactions to facilitate resolution, promoting M2 macrophage polarization, and stimulating angiogenesis, cell proliferation, and ECM remodeling ^47,50,51,58^. Indeed, we reason that the early transition to a NR-mediated regenerative response is part of the therapeutic mechanism achieved by human vascular progenitor cells delivered in infarcted hearts.

TGFβ has been shown to promote the NR phenotype ^56,68^. NRs in the injury site derive from the TGFβ-mediated polarization of naïve neutrophils (N0) recruited from the bone marrow (BM) ^68^. Blocking TGFβ signaling results in a depletion of the NR subpopulation and leads to a loss of vascularization ^47^. We have shown that human ECFC+MSC implanted in the subcutaneous space of immunodeficient mice can form robust vascular networks connected with the host circulation within a week ^42,48,72,73^. This vascularization process required host neutrophils expressing TGFβ receptor 2 (TGFBR2) ^47^. Human ECFC+MSC implanted in neutrophil-depleted animals (by sub-lethal radiation) completely failed to vascularize; however, an adoptive transfer of BM-derived neutrophils from unirradiated wild-type donor mice fully rescued vascularization ^47^. Of note, the adoptive transfer of BM-derived neutrophils lacking TGFBR2 (from *Tgfbr2*^−*/*−^ mice; *LysM-Cre*::*Tgfbr2/loxP*) failed to rescue vascularization, which underscored the critical role of TGFβ-TGFBR2 signaling in the generation of functional NRs ^47^. Here, we further confirmed that the successfully engrafted human ECFCs are a critical source of active NR-polarizing factors. We found that the conditional medium from ECFC+MSC coculture in a GelMA hydrogel strongly promoted NR polarization, evidenced by the up-regulation of NR signature genes, *Arg1, Vegfa*, and *Il4*, in mouse BM neutrophils. ECFC-derived TGFβ1 was up-regulated only in the presence of MSCs. Furthermore, when the endothelial-perivascular cell interaction was disrupted by blocking Notch-mediated signaling, ECFCs failed to up-regulate TGFβ1 expression. Taken together, our data suggest that the beneficial effect of delivering human vascular progenitor cells in GelMA hydrogel was partly due to promoting TGFβ1 secretion and engaging host NRs, resulting in a pro-regenerative microenvironment that favors post-MI cardiac healing.

Over the years, numerous preclinical studies and clinical trials have evidenced the challenges of treating heart diseases with cell injections ^29^. Systemic infusion of cells has been shown to be inefficient at delivering cells to the heart. It could even cause adverse effects when the injected cells accumulate in other organs, such as the lungs ^74^. For local intracoronary injections, the injected cells need to undergo transendothelial migration to reach the infarcted areas ^32^. Studies have shown the risks of forming plugs when transplanted cells pass through narrow capillaries, which have diameters similar to cells ^29,75^. Thus, those blocked cells become the barrier that prevents other cells from traveling from the coronary vessels to the ischemic zone.

As one of the most straightforward routes to deliver cells to the heart, the intramyocardial injection has been frequently applied in preclinical animal studies and clinical trials ^27,76–78^. With a needle or catheter, an intramyocardial injection can precisely deliver a sustainable number of therapeutic cells into pathological tissues in a short period of time. The intramyocardial injection can be performed by puncturing the myocardium from the epicardial side with a needle and injecting cells ^29^. The epicardial intramyocardial injection is common in small animal models and was used in this study. However, this method requires open-chest surgery to expose the heart, which limits its clinical application. The endocardial intramyocardial injection also requires penetration of the myocardium, but instead, from the inside of the ventricular wall ^78^. In human patients, this method can be achieved by a long catheter threaded through the peripheral blood vessels into the left ventricle, where the cells are injected through the endocardium. In general, this approach is less invasive because of it could avoid open-chest surgery ^79^. For this reason, the endocardial intramyocardial injection possesses a higher translational potential ^80^. However, it is also challenging to perform in small animal models.

A common hurdle is shared by all current delivery routes: low cell retention due to the “washout” effects of beating hearts. Given that the heart is a contractile pump, the beating of cardiomyocytes squeezes the narrow space in the myocardium and pushes transplanted cells in fluid medium out through the needle track or damaged blood vessels ^29^. Many studies had reported a rapid loss of >80% of injected cells within hours after delivery ^30,78^. Besides blood flow, the cells can be washed out of the heart by lymphatic drainage. In addition to physical washout from the heart, transplanted cells are lost over time through cell death before conferring a therapeutic benefit to the injured cardiac tissue. Inflammatory cells and damaged cardiomyocytes in the infarcted tissue can release cytokines and reactive oxygen species, which results in transplanted cell apoptosis ^81^. Furthermore, the infarcted microenvironment is deficient in oxygen and nutrient supply. As a result, many transplanted cells die in these unfavorable living conditions.

A myriad of studies confirms that cell retention plays a crucial role in the success of cell-mediated cardiac repair ^29^. Any delivery route needs to maintain donor cells in the recipient’s heart for enough time to send paracrine signals to surrounding damaged heart cells and repair them. Challenges associated with the in vivo cell delivery of currently available vehicles have limited clinical translation of this technology. To improve the retention of therapeutic cells in the heart, scientists have been focusing on bioengineering strategies, including designing novel biomaterials that protect cells from the washout effect in the heart ^7,82^.

Several injectable hydrogels have been designed with different base materials and cell type combinations ^36^. Previously, synthetic polymers and nanogels were used to encapsulate cells for delivery into mouse and swine models of MI to demonstrate their safety and efficacy ^37–39^. Notwithstanding the benefit of these new materials, the translational usage of synthetic materials faced obstacles due to the limited data on their long-term biocompatibility. Various natural extracellular matrix (ECM) products have been extensively tested for intramyocardial cell injection, including fibrin- or collagen-based hydrogels ^31^. However, the slow gelation process of these natural hydrogels failed to achieve desirable cell retention. Accordingly, improving the properties of naturally occurring ECMs has become an alternative approach. This can be achieved by the chemical functionalization of proteins, an approach that was proposed precisely to improve the usability of biomaterials ^83–86^. Here, we used GelMA synthesized by adding methacrylate groups to the amine-containing side-groups of gelatin, which becomes a photocrosslinkable, bioadhesive hydrogel ^40,41^. As a result, GelMA hydrogels exhibit characteristics of both natural and synthetic biomaterials. GelMA is highly soluble in phosphate-buffered saline and can be prepared as a precursor solution (5-20% w/v) at room temperature. Therapeutic cells can be easily resuspended in the GelMA precursor solution for needle- or catheter-based injections. Notably, the photocrosslinking reaction produces a GelMA hydrogel within 10 s of UV illumination without compromising encapsulated cell viability. Consequently, the quick gelation process of GelMA hydrogel effectively stops the washing out of cells injected into the hearts, resolving the main physical problem of low cell retention. Furthermore, GelMA hydrogel possesses excellent biocompatibility. In our previous studies, GelMA hydrogel provided an ideal microenvironment for the engraftment and vascularization of human vascular progenitor cells both in vitro and in vivo in immunodeficient mice ^41,42^. In particular, GelMA contains gelatin as its backbone, which provides cell-responsive characteristics such as the provision of appropriate cell adhesion sites and proteolytic degradability. Degraded products of GelMA hydrogel are simply gelatin peptides, which are parts of natural ECM components and are non-cytotoxic or non-immunogenic to the hearts ^40^.

The intensity needed to photocrosslink GelMA hydrogels transmyocardially was analyzed for safety. Transmyocardial exposure to our UV light spectrum (UVA-visible: 320-500 nm) had little observable effect on the human cells and surrounding mouse tissues ^42^. We tested these effects previously by exposing human cells and rodent skins to prolonged UV illumination (120 seconds) and showed minimal effects on cell apoptosis ^42^. In this study, we achieved sufficient transmyocardial GelMA photocrosslinking with only 10 s of UV illumination. Therefore, our working range of UV light exposure was deemed sufficiently safe. Alternatively, GelMA could be formulated to form a hydrogel with visible light-responsive photoinitiators to eliminate the usage of UV light ^87^, although the crosslinking time would be significantly longer than with UV light ^88^. Also, UV light illumination could be used through a fiber-optic system together with a catheter, as we previously showed that with a photocurable polymer patch to fix septal defects in the beating hearts of pigs ^89^. Patches attached to the heart by UV crosslinking showed similar inflammatory response as sutures, with 100% small animal survival, indicating safety and applicability of UV system in minimally invasive delivery in hearts.

In situ, the photocrosslinked GelMA hydrogel functions as a cell delivery vehicle in our study. We had shown previously that the implantation of GelMA hydrogel alone (containing no cell) provoked minimal host cell infiltration, suggesting GelMA is a relatively inert material ^41^. A recent study by Ptaszek et al. applied acellular GelMA hydrogel on top of the epicardial surface of the mouse heart at the time of experimental MI ^90^. Mice treated with these bioadhesive GelMA patches displayed improved post-MI survival rates and left ventricular contractile function, raising the opposability that our GelMA hydrogel may benefit cardiac healing by providing mechanical support. However, we compared our intramyocardial GelMA-cell injection to their GelMA bioadhesive application and found important differences between the two experimental conditions.

In our study, we used minimal UV exposure time, and thus the mechanical properties of GelMA hydrogel were relatively low compared to the GelMA bioadhesive protocol (elastin modulus, ∼2 kPa vs. ∼70 kPa). We reason that the disparity came from the fundamental difference in purpose between the two studies. We chose our GelMA crosslinking condition to optimize the viability and vasculogenic potential of the encapsulated human ECFCs and MSCs. We determined that a soft GelMA hydrogel (2 to 5 kPa) was permissive for the formation of functional vascular networks ^41^. In contrast, the study by Ptaszek et al.aimed to reinforce the loss of mechanical strength of the infarcted myocardial wall ^90^. They achieved a stiffer GelMA patch through a significantly longer (up to 4 min) UV illumination. For this purpose, we anticipated that our cell-laden GelMA hydrogel injected intramyocardially is unlikely to provide compatible mechanical support, as shown by Ptaszek et al.

Our study had several limitations, including the need to use immunodeficient mice to enable the transplantation of human cells. Nevertheless, although the SCID mice used in this study have a genetic immune deficiency that affects their B and T cells, their myeloid cell lineage is normal. Indeed, we did not notice dysregulated neutrophil behaviors in the absence of functional B and T immunity compared to other studies of neutrophil recruitment post-MI in immunocompetent mice ^50,52^. Also, we previously validated the dependency of graft vascularization on host neutrophils using fully immunocompetent hosts by transplanting mouse autologous endothelial cells and MSCs into syngeneic recipients ^47^. Therefore, we have demonstrated that neutrophil participation is critical in both syngeneic (immunocompetent recipients) and xenograft (immunodeficient recipients) cell engraftment models. In both cases, lymphocyte presence in the grafts was minimal. Nevertheless, the difference in the immune system between humans and rodents is well recognized, and thus our preclinical data will need to be further validated in human patients.

Another limitation of the mouse MI model is the difficulty performing more clinically relevant ischemic injuries. For instance, we induced MI by permeant ligation of LAD in mouse hearts, but a cardiac ischemia-reperfusion injury (IRI) would have been clinically more relevant ^26^. However, performing IRI in small mouse hearts is challenging due to significant variations and problems of reproducibility. Also, the most likely intramyocardial cell delivery route in the clinics would be through a minimally invasive catheter procedure to penetrate the endocardial side. However, it is extremely difficult to perform this procedure in mice and rats. Lastly, the analyses of cardiac functions in mice are limited to imaging-based measurements (echocardiography and magnetic resonance imaging) for live animals or histological examination for end-point experiments. Future studies in large animals could allow for more precise analysis, including the invasive hemodynamics with PV loops for long-term evaluations.

## 4. Conclusion

In summary, we have developed a novel therapeutic cell delivery method for MI treatment. Our approach entails the intramyocardial injection of vascular progenitor cells resuspended in a GelMA precursor solution followed by transmyocardial UV illumination, resulting in an in-situ photocrosslinked hydrogel that effectively retains the cells inside the ischemic myocardial tissue. This approach maintained high viability and cell retention, providing a significant advantage over cells injected in liquid or unmodified ECM gels. The improved cell retention allowed us to investigate the therapeutic effects of the human vascular progenitor cells post-MI. We demonstrated that the beneficial myocardial remodeling and stabilization of cardiac functions post-MI was enabled by the engrafted cells via the engagement and polarization of host pro-regenerative neutrophils through TGFβ signaling. This proof-of-concept study warrants further investigations, including scaling up for large animal tests, integrating a catheter-mediated injectate and a UV light delivery system, and potential modifications to include additional cell types such as iPSC-derived cardiomyocytes ^91^. We envision that our research provides an opportunity to generate an off-the-shelf cell-based therapy with the potential to treat acute myocardial infarction.

## 5. Experimental Section

### Cell culture

Human ECFCs and MSCs were isolated from human umbilical cord blood and subcutaneous adipose tissue, respectively, as previously described ^45,92^. ECFCs were cultured on 1% (w/v) gelatin-coated plates and maintained in ECFC-medium: EGM-2 (except for hydrocortisone; PromoCell) supplemented with 20% FBS (Hyclone) and 1X glutamine-penicillin-streptomycin (GPS; Invitrogen). ECFCs express CD31, VE-cadherin, von Willebrand factor (vWF), but not CD90, CD45 or CD14. MSCs were cultured on uncoated plates using mesenchymal stem cell growth medium (MSCGM; ATCC) with MSC growth supplement (ATCC) and 1X GPS. MSCs express CD90 and PDGFRβ, but not endothelial or hematopoietic markers. ECFCs and MSCs between passages 6 and 12 were used for all the experiments.

### Synthesis of GelMA

GelMA was synthesized as previously described ^41^. Briefly, porcine skin gelatin (type A; Sigma-Aldrich) was dissolved in phosphate-buffered saline (PBS) at 60 °C to make a 10% (w/v) gelatin solution. Methacrylic anhydride (Sigma-Aldrich) was slowly added to the gelatin solution under stirring conditions to a final concentration of 1% (v/v). The mixture was allowed to react for 3 h at 50 °C. The product was dialyzed against deionized water to remove unreacted methacrylic anhydride. The dialyzed GelMA solutions were lyophilized and stored at -80 °C. The degree of methacrylation was quantified by an NMR spectrometer (Varian INOVA) to be 49.8% functionalized to original amino groups.

### Preparation of GelMA precursor solution

A GelMA precursor solution was prepared by dissolving lyophilized GelMA (5 w/v% final) and photoinitiator Irgacure 2959 (0.5 w/v%) in PBS at 80 °C and then cooled to 37 °C in a water bath. Human ECFCs and MSCs (5×10^5^ cells for each cell type) were resuspended in 100 μL of GelMA precursor solution. The cell/GelMA mixture was kept at 37 °C, protected from light, and used within 1 h.

### Mouse model of myocardial infarction and intramyocardial cell delivery

All animal experiments were conducted under a protocol approved by the Institutional Animal Care and Use Committee at Boston Children’s Hospital. NOD.SCID mice (NOD.Cg-*Prkdc*^scid^/J; male, aged 10-12 weeks, weighing 30-35 g; Jackson Laboratory) were anesthetized using an isoflurane chamber (1-3% isoflurane) and given pre-operative pain management. The mouse tongue was retracted and held with forceps, and a 20G catheter was used to intubate into the trachea. The catheter was then attached to the small animal ventilator. Ventilation was performed with a tidal volume of 200 μl and a respiratory rate of 133/min. 100% oxygen was provided to the inflow of the ventilator. Following anesthesia, the mice were fixed on a heating pad to maintain normothermia. The anterior surface of the heart was exposed by a left thoracotomy between the third and fourth ribs. MI was induced by the permanent ligation of the left anterior descending (LAD) artery with an 8-0 nylon suture. Criteria for occlusion success was an acute color change in the left ventricular wall (red to pale). Afterward, the cell/GelMA mixture was injected using a 30G needle into the myocardium and followed by UV photocrosslinking. OmniCure S2000 UV lamp (Lumen Dynamics) was used to provide UV exposure in this study. UV intensity was adjusted using a UV intensity meter (G&R Labs). In situ polymerization of cell-laden GelMA hydrogel was achieved by transmyocardial exposure to UV light (intensity of 40 mW/cm^2^) for 10 seconds. The thorax was closed with absorbable sutures. The air was withdrawn from the chest using a syringe with a 26G needle. Following the recovery of spontaneous breathing, the mice were extubated and returned to its cage.

### Quantification of cell retention by in vivo bioluminescence imaging

Luciferase-expressing ECFCs (luc-ECFCs) were generated by transfection of a PiggyBac vector carrying a CMV promoter-driven firefly luciferase reporter gene with a super PiggyBac transposase expression vector (System Biosciences) ^93^. The ratio between transposon and transposase vectors was 5:1, and a total of 2.4 μg of DNA was used to transfect 1×10^6^ ECFCs. After puromycin selection, this transposon system achieved a stable expression of the luciferase reporter gene in ECFCs. Luciferase-expressing ECFCs were suspended in 100 μL of GelMA precursor solution in the presence of non-labeled MSCs (Luciferase-expressing ECFC: MSC = 1:1; total 1×10^6^ cells). The cell suspension was injected into the infarcted myocardium followed by UV illumination (GelMA-H group). Cell injection without UV illumination (GelMA-S group) served as a control. On 3, 24, 48, and 72 hours after cell injections, the entire body of each mouse was imaged using an IVIS 200 Imaging System (Xenogen Corporation). Briefly, mice were anesthetized using an isofluorane chamber and were given the substrate, luciferin (Promega), by intraperitoneal injection according to body weight (125 mg/kg). Bioluminescence was detected for 5 min after luciferin administration, and the collected data was analyzed with Live Image 3.0 (Xenogen Corporation). The same cell suspension injected into arrested mouse hearts and performed a bioluminescence measurement after 15 min served as a baseline.

### Echocardiographic measurement

Cardiac function following LAD ligation and treatments were evaluated by using a high-frequency ultrasound system Vevo 2100 (VisualSonics) with a 30-MHz central frequency scan head. To facilitate the echocardiography, the mice were anesthetized with isoflurane and placed on a heated pad in the left lateral decubitus position. Two-dimensional echocardiography and M-mode images were obtained using a short axis view from the mid-LV at the tips of the papillary muscles. The left ventricular ejection fraction (LVEF) and left ventricular fractional shortening (LVFS) were calculated from LV dimensions in the 2D short axis view. Echocardiography was performed on each mouse 1 h before LAD ligation to obtain a reference of baseline cardiac function. Cardiac function was monitored post-MI at 3 days and 1, 2, and 4 weeks.

### Histology and immunofluorescence staining

At indicated time points, the mouse hearts were harvested and weighed, fixed in 10% neutral-buffered formalin overnight, and embedded in paraffin. Each paraffin-embedded heart was sectioned (7 μm thick) through the infarcted area and stained with Hematoxylin-eosin or Masson’s trichrome staining. Myocardial wall thickness and fibrosis were calculated by computerized planimetry using ImageJ software, version 1.44 (NIH). For immunostainings, sections were deparaffinized and rehydrated through sequential immersion in xylene, 100%, 90%, 80% and 50% ethanol. Antigen retrieval was carried out by heating the sections in Tris-EDTA buffer at 95 °C for 30 min. Sections were then blocked for 30 min in 5% blocking serum followed by incubation with primary antibodies overnight at 4 °C. Human-specific vimentin antibody (clone V9; Abcam) and the lectin Ulex Europaeus Agglutinin I (UEA-I; Vector Laboratories) were used to stain human blood vessels ^93^. Fluorescent secondary antibodies were applied to the sections for 1 h at room temperature, followed by nuclei counterstaining with DAPI. Finally, sections were mounted with a fluorescent mounting medium (Dako).

### Flow cytometry

Mouse hearts were harvested from euthanized mice and enzymatically digested with collagenase A (1 mg/mL; Roche Life Science) and dispase (2.5 U/mL; BD Biosciences) for 2 h at 37 °C. The retrieved cells were incubated with PerCP-conjugated anti-mouse CD45 (1:100; BD Biosciences), FITC-conjugated mouse Ly6G (1:100; clone 1A8, Biolegend), PE-conjugated anti-mouse F4/80 (1:100; eBiosciences), APC-conjugated anti-mouse CD206 (1:50; Biolegend), and APC.Cy7-conjugated anti-mouse CXCR2 (1:50; Biolegend) antibodies. Flow cytometric analyses were performed using a BD LSRFortess flow cytometer (BD Biosciences) and FlowJo software (Tree Star Inc.). Murine hematopoietic cells were identified as mCD45^+^ cells. Ly6G^+^F4/80^-^ and Ly6G^-^ F4/80^+^ mouse hematopoietic cells were neutrophils or monocytes/macrophages, respectively. Within Ly6G^+^F4/80^-^ neutrophils, neutrophil subpopulations were classified into naïve neutrophils (N0; CXCR2^-^), pro-inflammatory neutrophils (NI; CXCR2^+^CD206^-^), and pro-regenerative neutrophils (NR; CXCR2^+^CD206^+^).

To determine human ECFC and MSC retention, the retrieved cells were stained with PerCP-conjugated anti-mouse CD45, FITC-conjugated anti-human CD31 (1:50 Biolegend), and PE-conjugated anti-human CD90 (1:100 Biolegend). ECFCs were identified as mCD45^-^ hCD31^+^hCD90^-^ cells and MSCs were mCD45^-^hCD31^-^hCD90^+^.

### RNA-sequencing (RNAseq) analysis

RNAseq was performed on infarcted myocardial tissues (GelMA-H and GelMA-S groups) on day 2 post-MI. Tissues from uninjured hearts served as controls. Each group consisted of 3 biological replicates. Total RNA was extracted using RNeasy Mini Kit (Qiagen) following the manufacturer’s protocol. RNA quantity and quality were checked with Agilent Bioanalyzer instrument. Libraries were prepared and sequenced by GENEWIZ (NJ, USA). Library preparation involved mRNA enrichment and fragmentation, chemical fragmentation, first and second strand cDNA synthesis, end repair and 5’ phosphorylation, Da-tailing, adaptor ligation, and PCR enrichment. Libraries were then sequenced using Illumina HiSeq2500 platform (Illumina) using 2×150 paired end configuration. Raw sequencing data (FASTQ files) was examined for library generation and sequencing using FastQC (Babraham Institute) to ensure data quality. Reads were aligned to UCSC mm10 genome using the STAR aligner ^94^. Alignments were checked for evenness of coverage, rRNA content, genomic context of alignments, complexity, and other quality checks using a combination of FastQC and Qualimap ^95^. The expression of the transcripts was quantified against the Ensembl release GRCm38 transcriptome annotation using Salmon. These transcript abundances were then imported into R (version 3.5.1) and aggregated to the gene level with tximport. Differential expression at the gene level was called with DESeq2 ^96^. Pairwise differential expression analysis between groups was performed using Wald significance test. *P* values were corrected for multiple hypothesis testing with the Benjamini-Hochberg false-discovery rate procedure (adjusted *P* value). Genes with an adjusted *P*-value < 0.05 and absolute log2 fold change > 1 were called as differentially expressed genes. PCA analysis was performed on DESeq2 normalized, rlog variance stabilized reads. All samples comparison was performed using Likelihood Ratio Test (LRT). Functional enrichment of differential expressed genes, using gene sets from GO, was determined with Fisher’s exact test as implemented in the cluster Profiler package. Heat maps of differential expressed genes and enriched gene sets were generated with pheatmap package. Volcano plots of enriched gene sets were generated with ggplot2 package.

### Quantitative Real-Time PCR analysis

Total RNA was extracted from cells using the RNeasy Mini Kit (Qiagen) following the manufacturer’s workflow. RNA concentration was measured by the NanoDrop 8000 spectrophotometer (Thermo Fisher), and RNA purity was evaluated by the ratio of absorbance at 260 and 280 nm. The cDNA synthesis was processed by High-Capacity RNA-to-cDNA Kit (Thermo Fisher). The quantitative real-time PCRs were performed on the QuantStudio 6 Flex Feal-Time PCR System with PowerUp SYBM Green Master Mix (Thermo Fisher). GAPDH served as the housekeeping gene. Sequences of primers for real-time PCR are listed in Table S1, Supporting Information.

### ECFC-MSC coculture and conditioned medium generation

Samples of conditioned medium were collected from GelMA hydrogels containing ECFC+MSC coculture (1:1 ratio; total 5×10^5^ cells per 100 μL gel) or monoculture (only ECFC or MSC alone; total 5×10^5^ cells per 100 μL gel) over eight days in vitro. To this end, cell-laden GelMA hydrogels were cultured in 3-ml tubes with 500 μL of Basal medium (EBM-2, 5% FBS medium) refreshed every 24 h. Collected samples of conditioned medium were filtered (0.2 μm) and then concentrated ten-fold (Amicon Ultra centrifugal filters; 3 kDa cut off; Millipore).

To determine the effect of ECFC-MSC coculture on TGFβ1 expression, cells were retrieved from GelMA hydrogels after in vitro coculture for 8 days by enzymatical digestion (collagenase/dispase; 1h at 37 °C). ECFCs were sorted out from MSCs by MACS using magnetic beads (Dynabeads; Thermo Fisher Scientific) coated with anti-human CD31 antibodies. The purity of MACS-sorted cells was validated by the cell-type specific mRNA expressions of human *CD31* or *PDGFRβ* for ECFCs or MSCs, respectively, by the quantitative real-time PCR analysis. Retrieved cells from gels containing only ECFC or MSC alone served as controls. For Notch-signaling inhibition studies, GelMA hydrogel containing ECFC+MSC coculture were treated with the γ-secretase inhibitor DATP (10 μM in EGM-2 with 5% FBS; Selleckchem) for 24 h before harvesting conditioned media ^47^.

### Human cytokine protein array

The presence of selected cytokines was evaluated in samples of conditioned medium with Proteome profiler human angiogenesis array (R&D Systems) according to the manufacturer’s instructions. Antigen-antibody reactions were visualized using LumiGLO substrate (Kirkegaard & Perry Laboratories, Inc.) and chemiluminescent sensitive film (Kodak). Densitometry was performed by image analysis (ImageJ) to estimate the amount of protein present in each sample.

### Isolation of mouse bone marrow neutrophils

Mouse femur bone was dissected from euthanized mice and cut at both ends. A 23G needle was used for flushing the bone marrow out with ice-cold HBSS buffer. After RBC lysis, mouse bone marrow neutrophils were sorted from the bone marrow cell suspension using a mouse neutrophil isolation kit that enriches for CD11b^+^Ly6G^+^ cells (Miltenyi Biotec; negative selection using a cocktail that contains anti-CD5, -CD45R, -CD49b, -CD117, -F4/80 and -Ter119 antibodies). Isolated neutrophils were maintained in StemSpan H3000 medium (STEMCELL Technologies) supplemented with GlutaMAX (1X; Thermo Fisher Scientific), ExCyte (0.2%; Merck Millipore), Am580 retinoic acid agonist (2.5 mM; Sigma-Aldrich) and human G-CSF (150 ng/mL; PeproTech).

### In vitro neutrophil polarization assay

Due to the limited lifespan of neutrophils, the freshly isolated mouse bone marrow neutrophils were used immediately for conditioned medium studies. Mouse bone marrow neutrophils (5×10^6^ cells) were cultured for 24 h in 1 mL of StemSpan H3000 medium supplemented with ten-fold concentrated conditioned media (reconstituted in 9:1 volume ratio). Basal medium (EBM-2, 5% FBS medium) served as a control. After 24 h, neutrophils were collected for quantitative PCR analysis of NR polarization markers (*Il4, Vegfa*, and *Arg1*). Mouse neutrophils treated with recombinant human TGFβ1 (10 ng/mL; PeproTech) for 24 h served as a positive control for NR polarization ^56^.

### Neutrophil depletion

For neutrophil depletion studies, either anti-mouse Ly-6G/Ly-6C Gr1 (Bio X Cell) or control (IgG_2b_; Bio X Cell) antibodies were administered intraperitoneally into mice every 2 days from 2 days before LAD ligation to post-operative day 6. Anti-Gr1 given at 200 μg/mouse was shown to be sufficient to deplete neutrophils in the circulation, confirmed by flow cytometry using FITC-conjugated Ly6G antibody (1:100; clone 1A8, Biolegend) ^46^.

### Microscopy

Images were taken by the Axio Observer Z1 inverted microscope (Carl Zeiss) and AxioVision Rel.4.8 software. Fluorescent images were taken with an ApoTome.2 Optical sectioning system (Carl Zeiss) and 20X or 40X objective lens. Non-fluorescent images were taken with an AxioCam MRc5 camera using a 10X or 20X objective lens.

### Statistical analyses

All statistical analyses were performed using the GraphPad Prism v.7 software (GraphPad Software Inc.). The sample size, including the number of mice per group, was chosen to ensure adequate power and based on historical laboratory data. No exclusion criteria were applied for all analyses. All data were expressed as mean ± standard deviation of the mean (s.d.). Comparisons between multiple groups were performed by ANOVA followed by Bonferroni’s post-test analysis. Unpaired two-tailed Student’s t-test was used for comparisons between two groups. A value of *P*<0.05 was considered to be statistically significant.

## Acknowledgments

This work was supported by grants from the National Institutes of Health (R01AR069038 and R01HL128452 to J.M.M.-M.).

## Author contributions

XH, JMM-M and R-ZL conceived and designed the project. XH, ACL, ID, NO, G-BI, PdN-J, JMM-M, and R-ZL performed the experimental work. All authors discussed and analyzed the data and edited the results. XH, JMM-M and R-ZL wrote the manuscript.

## Data availability

The authors declare that all data supporting the findings of this study are available in the paper and its Supplementary Information.

