## Supplemental figures for "A photopolymerizable hydrogel enhances intramyocardial vascular cell delivery and promotes post-myocardial infarction healing by polarizing pro-regenerative neutrophils"

### Supplemental Tables

#### Table S1 – Sequences of primers used for Quantitative Real-Time PCR analysis

| Species* | Gene | Forward (5’ to 3’) | Reverse (5’ to 3’) |
|---|---|---|---|
| **h** | CD31 | ATTGCAGTGGTTATCATCGGAGTG | CTCGTTGTTGGAGTTCAGAAGTGG |
| **h** | PDGFRβ | AGCACCTTCGTTCTGACCTG | TATTCTCCCGTGTCTAGCCCA |
| **h** | TGFβ1 | CCCAGCATCTGCAAAGCTC | GTCAATGTACAGCTGCCGCA |
| **h** | GAPDH | AGGGCTGCTTTTAACTCTGGT | CCCCACTTGATTTTGGAGGGA |
| **m** | Il4 | GAGTCCAAGTCCACATCACTGAA | TGGCTCAGTACTACGAGTAATC |
| **m** | Vegfa | GGAAAGACCGATTAACCATGTCA | GGCTTTCTGGATTAAGGACTGTTC |
| **m** | Arg1 | CCACAGTCTGGCAGTTGGAA | GCATCCACCCAAATGACACA |
| **m** | Gapdh | CATTTGCAGTGGCAAAGTGGAG | CAATCTTGAGTGAGTTGTCATAT |

* Primers were designed to specifically recognize human (h) or mouse (m) genes.

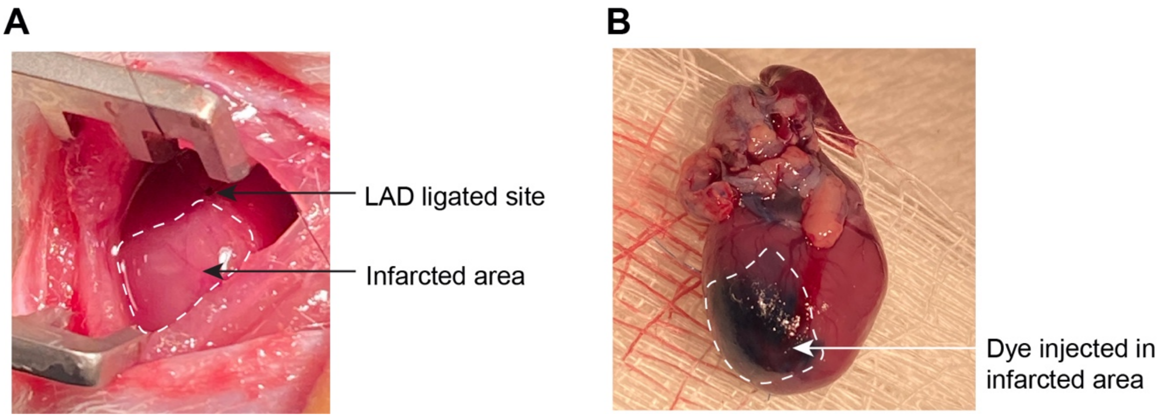


**Figure S1. Representative images of surgically induced myocardial infarction in mice**. (A) The left ventricle region of a mouse heart after the ligation of the left anterior descending (LAD) artery. The blenching region (marked by a white dashed line) indicated the infarcted area. (B) To determine the distribution of injectate in the infarcted heart, we added trypan blue in our GelMA precursor solution and injected a total volume of 100 μL per heart. The dye-labeled area indicated that the injection volume was sufficient to cover the infarcted zone.

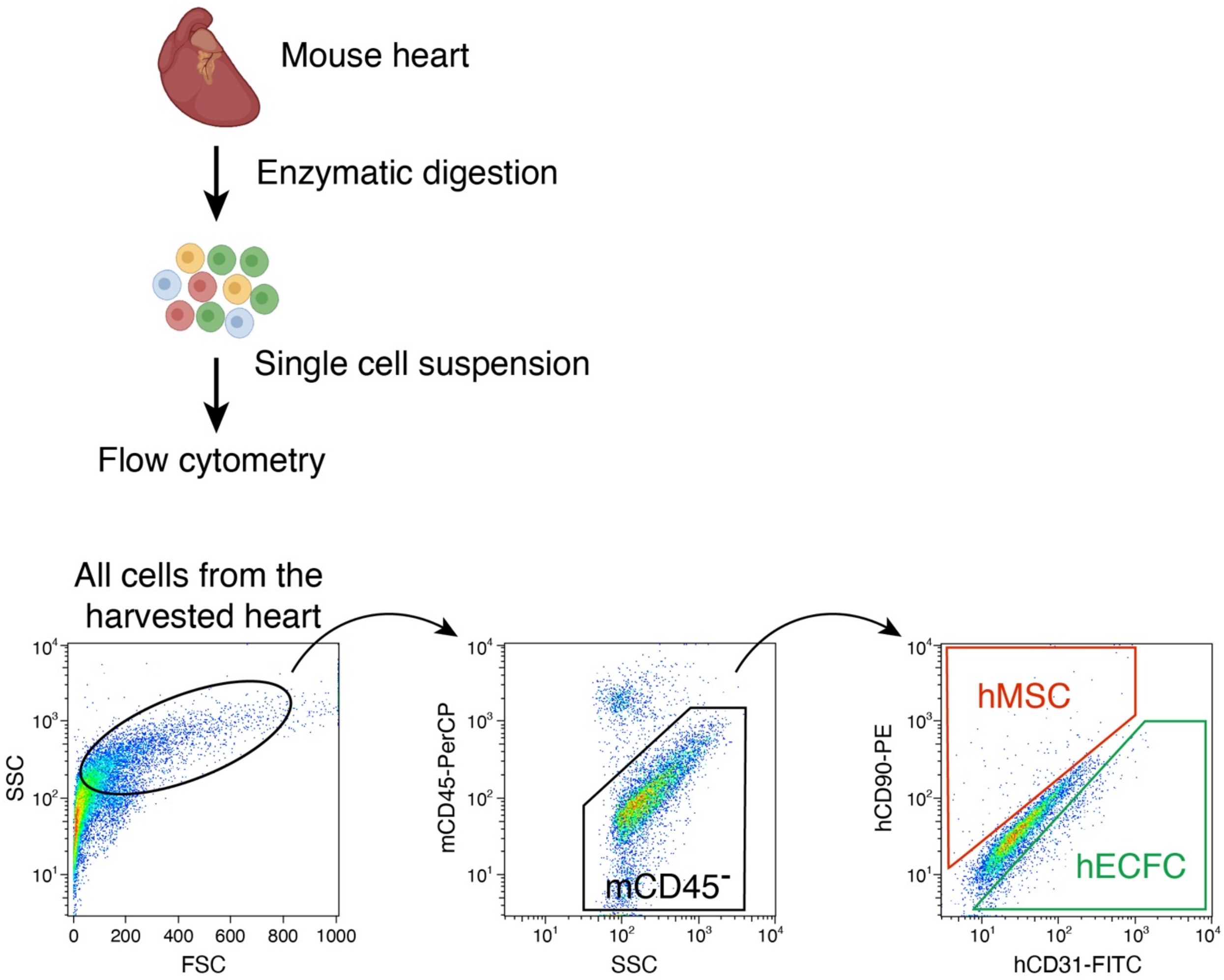


**Figure S2. Flow cytometry analysis of human cell retention in infarcted hearts.** General scheme for flow cytometry analysis of human ECFCs and MSCs delivered in GelMA-H at 48 h post-MI. Single-cell suspension was obtained by enzymatic digestion of harvested mouse hearts. Non-mouse hematopoietic cells were first gated by the negative fraction of PerCP-conjugated anti-mouse CD45 antibodies. Then, the human ECFCs and MSCs were detected by the FITC-conjugated anti-human CD31 and PE-conjugated anti-human CD90 antibodies, respectively.

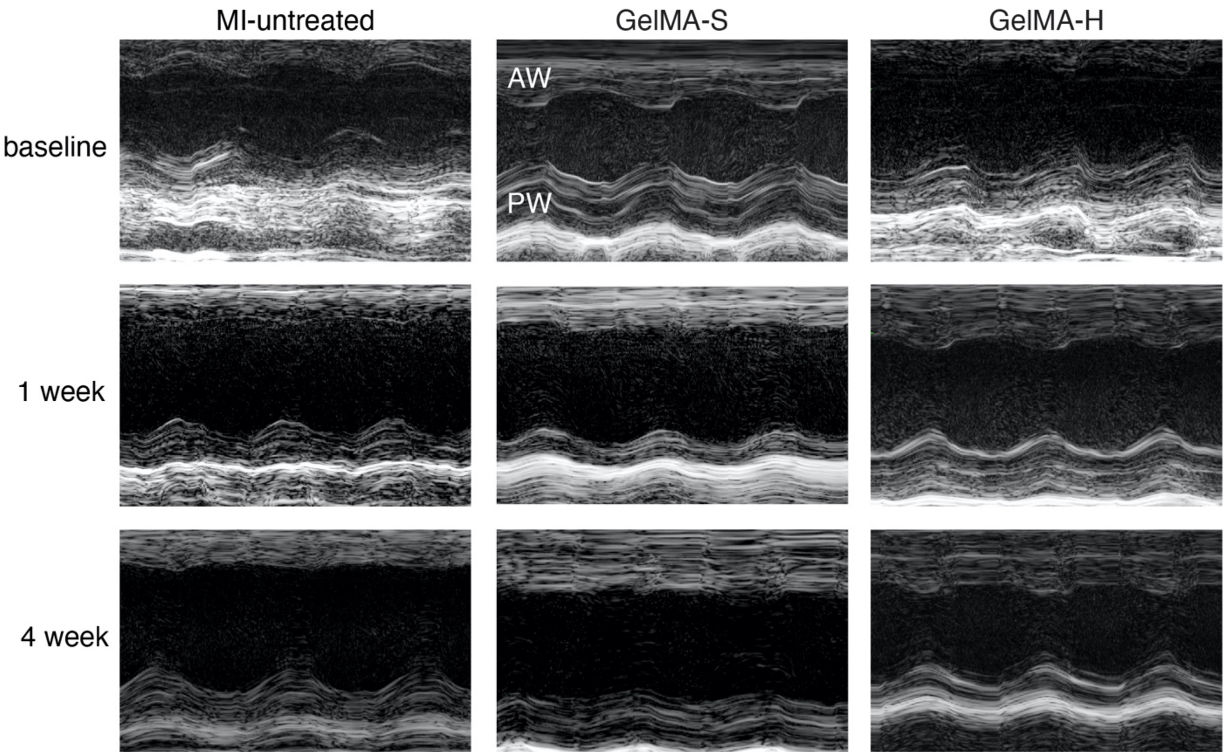


**Figure S3. Representative echocardiographic images (M-mode).** Each animal was measured 1 h before LAD ligation to obtain the baseline cardiac function. After MI induction, animals were divided into three groups: no cell injection (MI-untreated), ECFC+MSC injected in GelMA precursor solution (GelMA-S), and ECFC+MSC delivered in GelMA hydrogel (GelMA-H). Echocardiography was performed after 3 days, 1, 2, and 4 weeks post-MI. AW, anterior wall; PW, posterior wall.

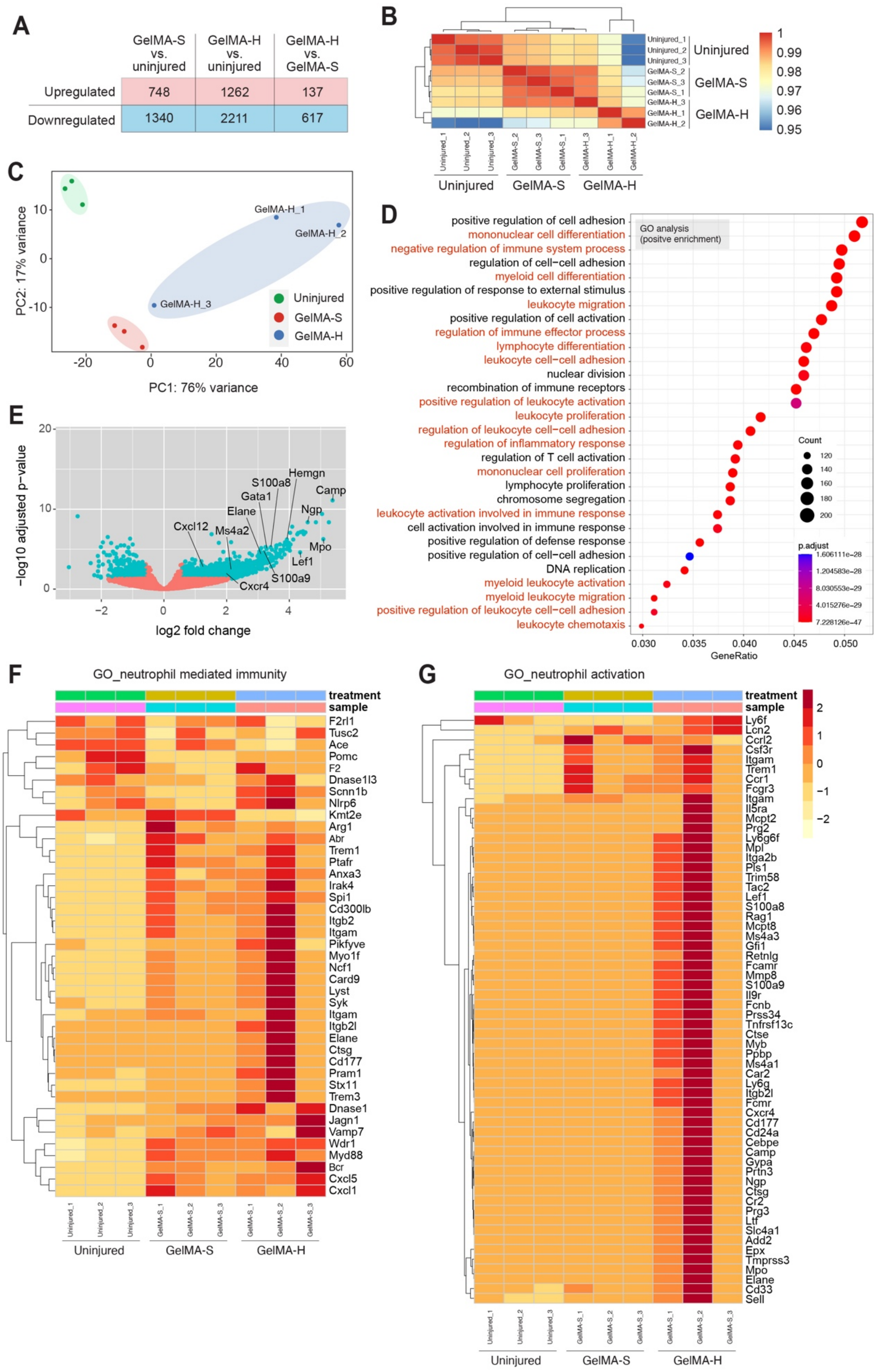
A
GelMA-S vs. uninjured
GelMA-H vs. uninjured
GelMA-H vs. GelMA-S
Upregulated 748 1262 137
Downregulated 1340 2211 617
B
Uninjured
GelMA-S
GelMA-H
1
0.99
0.98
0.97
0.96
0.95
C
PC1: 76% variance
PC2: 17% variance
GelMA-H_1
GelMA-H_2
GelMA-H_3
Uninjured
GelMA-S
GelMA-H
D
GO analysis
(positve enrichment)
positive regulation of cell adhesion
mononuclear cell differentiation
negative regulation of immune system process
regulation of cell−cell adhesion
myeloid cell differentiation
positive regulation of response to external stimulus
leukocyte migration
positive regulation of cell activation
regulation of immune effector process
lymphocyte differentiation
leukocyte cell−cell adhesion
nuclear division
recombination of immune receptors
positive regulation of leukocyte activation
leukocyte proliferation
regulation of leukocyte cell−cell adhesion
regulation of inflammatory response
regulation of T cell activation
mononuclear cell proliferation
lymphocyte proliferation
chromosome segregation
leukocyte activation involved in immune response
cell activation involved in immune response
positive regulation of defense response
positive regulation of cell−cell adhesion
DNA replication
myeloid leukocyte activation
myeloid leukocyte migration
positive regulation of leukocyte cell−cell adhesion
leukocyte chemotaxis
Count
120
140
160
180
200
p.adjust
GeneRatio
E
−log10 adjusted p−value
log2 fold change
Hemgn
S100a8
Camp
Gata1
Elane
Ngp
Cxcl12
Ms4a2
Mpo
Lef1
S100a9
Cxcr4
F
GO_neutrophil mediated immunity
treatment
sample
F2rl1
Tusc2
Ace
Pomc
F2
Dnase1l3
Scnn1b
Nlrp6
Kmt2e
Arg1
Abr
Trem1
Ptafr
Anxa3
Irak4
Spi1
Cd300lb
Itgb2
Itgam
Pikfyve
Myo1f
Ncf1
Card9
Lyst
Syk
Itgam
Itgb2l
Elane
Ctsg
Cd177
Pram1
Stx11
Trem3
Dnase1
Jagn1
Vamp7
Wdr1
Myd88
Bcr
Cxcl5
Cxcl1
Uninjured
GelMA-S
GelMA-H
G
GO_neutrophil activation
treatment
sample
Ly6f
Lcn2
Ccrl2
Csf3r
Itgam
Trem1
Ccr1
Fcgr3
Itgam
Il5ra
Mcpt2
Prg2
Ly6g6f
Mpl
Itga2b
Pls1
Trim58
Tac2
Lef1
S100a8
Rag1
Mcpt8
Ms4a3
Gfi1
Retnlg
Fcamr
Mmp8
S100a9
Il9r
Fcnb
Prss34
Tnfrsf13c
Ctse
Myb
Ppbp
Ms4a1
Car2
Ly6g
Itgb2l
Fcmr
Cxcr4
Cd177
Cd24a
Cebpe
Camp
Gypa
Prtn3
Ngp
Ctsg
Cr2
Prg3
Ltf
Slc4a1
Add2
Epx
Tmprss3
Mpo
Elane
Cd33
Sell
Uninjured
GelMA-S
GelMA-H

**Figure S4. Gene expression analysis following human vascular cells delivery in infarcted hearts.** Human ECFC+MSC were delivered in the GelMA precursor solution (Group: GelMA-S) or photocrosslinked GelMA hydrogel (Group: GelMA-H) in MI hearts. The infarcted left ventricle tissues were harvested on day 2 post-MI for RNAseq gene expression profiling. The uninjured hearts (Group: Uninjured) served as controls. (A) The number of differentially expressed genes (DEGs) between the three groups. (B) Hierarchical clustering analysis between all samples. (C) Principal components analysis (PCA). (D) Gene ontology (GO) analysis between GelMA-H and Uninjured groups showing the pathways related to immune response and neutrophil recruitment. (E) Volcano plot with up- and down-regulated genes comparing the GelMA-H and GelMA-S groups. (F-G) Heat maps and hierarchical clustering analyses of selected neutrophil-related genes between the three groups. Color bar indicates gene expression in scale.
